# A Transformer-Based Approach to Survival Outcome Prediction

**DOI:** 10.1101/2024.11.03.621674

**Authors:** Ted Mellors, Matt Schneider, Mehran Spitmann

**Author notes:** Conflicts of Interest: No conflict of interest.

## Abstract

Accurate prediction of patient survival outcomes is a critical challenge in cancer research, with the potential to inform personalized treatment strategies and improve patient care. We leveraged Geneformer, a state-of-the-art transformer model pre-trained on a massive single-cell RNA-seq dataset, to develop a model for the prediction of overall survival (OS). We adapted Geneformer for bulk tumor data analysis by appending a task-specific transformer layer and fine-tuning the model on RNA-seq data from The Cancer Genome Atlas (TCGA). Additionally, we employed a rank-value encoding scheme to prioritize informative genes and reduce noise. Our model demonstrated a robust correlation between predicted and true OS, with Pearson correlation coefficient of 0.72 (p<0.00001). Survival analysis revealed significant differences in survival between patient subgroups stratified based on the model’s predictions. The Geneformer-based model outperformed traditional machine learning approaches (Random Forest and Neural Network) in patient stratification tasks. Further analysis demonstrated the consistency of the model’s performance across different tumor stages and patient subgroups. Our study highlights the potential of leveraging pre-trained transformer models, originally developed for single-cell data analysis, to predict clinically relevant outcomes from bulk tumor gene expression data. The superior performance of our Geneformer-based model underscores its potential to enhance prognostication and treatment decision-making in cancer research. Future work will focus on refining the model architecture, incorporating multi-omics data, and validating its performance on external datasets to further advance its clinical utility.

**Short Abstract:** Accurate prediction of patient survival has important implications for cancer research as it enables the development of personalized treatment plans, guides clinical decision-making, and can be leveraged for clinical trial optimization. We utilized Geneformer, a transformer model pre-trained on single-cell RNA-seq data, to predict overall survival (OS) from bulk tumor gene expression. Adapting Geneformer for bulk tumor analysis and using rank-value encoding, we achieved strong correlations between predicted and true OS (r=0.72, p<0.00001). Our model outperformed traditional machine learning approaches in patient stratification, demonstrating consistent performance across tumor stages and subgroups. This study highlights the potential of pre-trained transformer models for prognostication in cancer, paving the way for refined, personalized treatment strategies.

## Introduction

The advent of high-throughput single-cell RNA sequencing (scRNA-seq) has revolutionized our understanding of cellular heterogeneity and its role in complex biological processes [1]. By providing a comprehensive snapshot of gene expression at the single-cell level, scRNA-seq enables researchers to unravel the intricate dynamics of gene regulatory networks and cellular states [2]. However, the sheer volume and complexity of scRNA-seq data present significant challenges in extracting meaningful insights. Traditional computational methods often struggle to capture the entire spectrum of gene expression patterns, particularly in the context of rare cell types or transient cellular states [3].

Recent advancements in deep learning, particularly transformer-based models, have shown immense promise in tackling the challenges posed by scRNA-seq data analysis [4]. These models, empowered by their capacity to capture long-range dependencies and contextual information, have demonstrated remarkable performance in tasks such as cell type identification, gene expression prediction, and trajectory inference [5, 6]. Building on these successes, we sought to leverage the power of transformer models to address a critical clinical challenge: the prediction of patient survival outcomes based on gene expression data from bulk tumor samples.

In this study, we utilized Geneformer, a state-of-the-art transformer model pre-trained on a massive single-cell RNA-seq dataset (Genecorpus-30M) [7], to develop a predictive model for the prediction of overall survival (OS). While Geneformer has demonstrated outstanding performance in single-cell gene expression prediction and classification tasks [7], its application to bulk tumor data and survival outcome prediction remains largely unexplored. To adapt Geneformer for our specific aims, we implemented key modifications to the model architecture and fine-tuning process. Specifically, we appended a task-specific transformer layer to the pre-trained Geneformer model and fine-tuned the model on bulk tumor RNA-seq data from The Cancer Genome Atlas (TCGA), with the objective of predicting OS. Additionally, we employed a rank-value encoding scheme to prioritize informative genes and reduce noise in the input data [7].

To thoroughly evaluate the performance of our models, we curated a cohort of patients from the TCGA dataset. The subsequent section details the patient selection process and the demographic and clinical characteristics of the included patients, providing critical context for the interpretation of our results.

## Results

### Patient Cohort and Data Characteristics

To assess the predictive capabilities of our models, we curated a patient cohort from The Cancer Genome Atlas (TCGA) dataset. Patients were included if they had available gene expression data, primary tumor samples, and documented records for either Days to Death (DTD) or Days to Last Follow-up (DTLF). Overall survival (OS) was defined as the number of days to death for patients with available DTD data; for patients without a recorded DTD, OS was determined using DTLF. To ensure adequate statistical power, we focused on resection sites and histologies with a minimum of 300 patients with OS data, including at least 25 OS patients with observed OS events (DTD). This resulted in a total of 3,254 patient samples evaluated in this study.

A complete overview of selected patient demographics and clinical characteristics can be found in **Table 1**. This table provides a detailed summary of key patient attributes, including age, gender, tumor stage, and other relevant clinical factors.

**Table 1.**
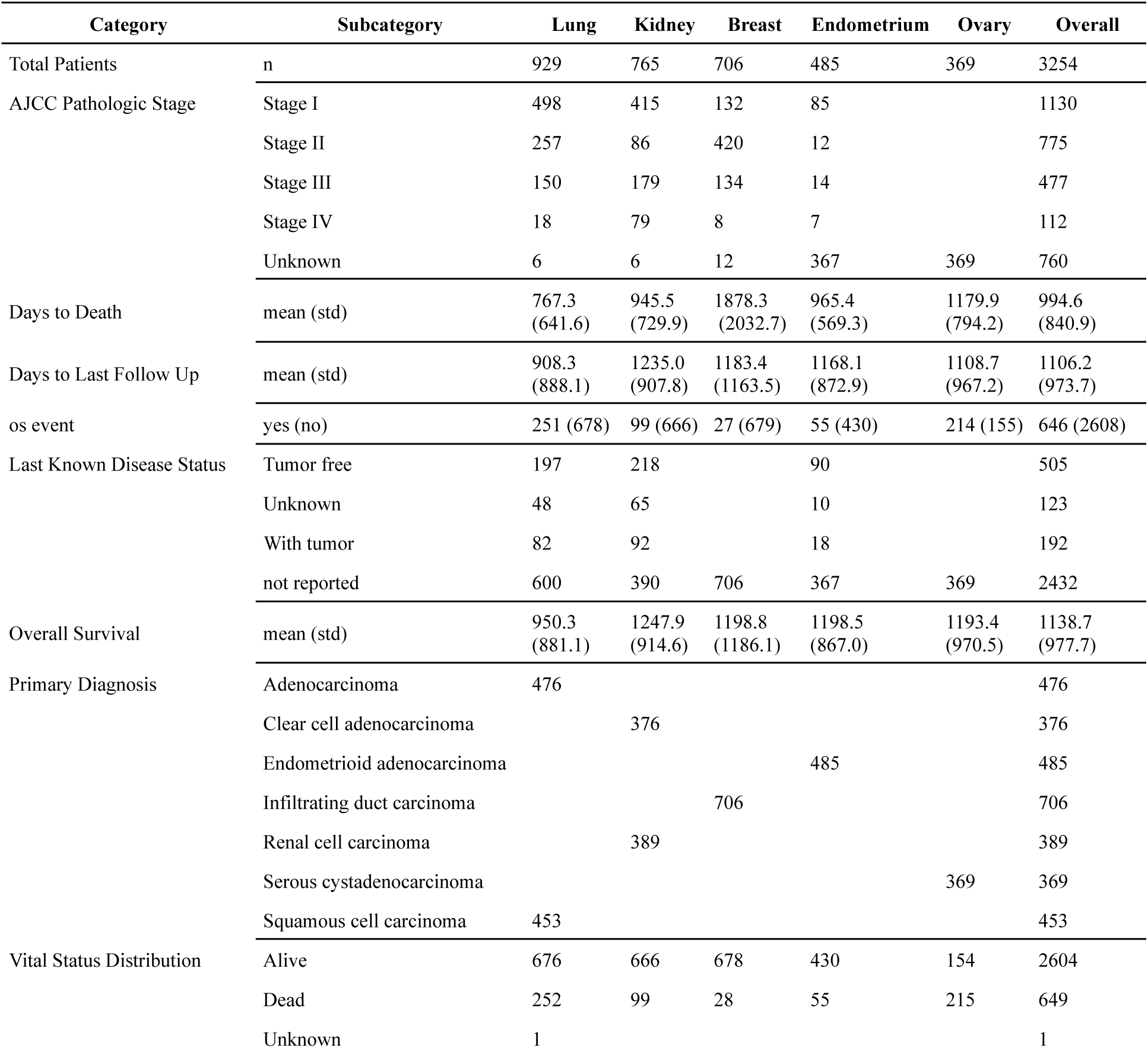
Demographics of study cohort selected for analysis from TCGA GDC Data Portal.

### Model Predictions and their Correlation with Clinical Outcomes

The model’s predictive capabilities were evaluated by assessing the correlation between the predicted values and the corresponding true OS values for the entire patient cohort (**Figure 1a**). A Pearson correlation of r = 0.72 (p<0.00001) was observed across all cancer types, indicating a substantial degree of overall concordance between predicted and true values regardless of tumor resection site or histology.

**Figure 1.**
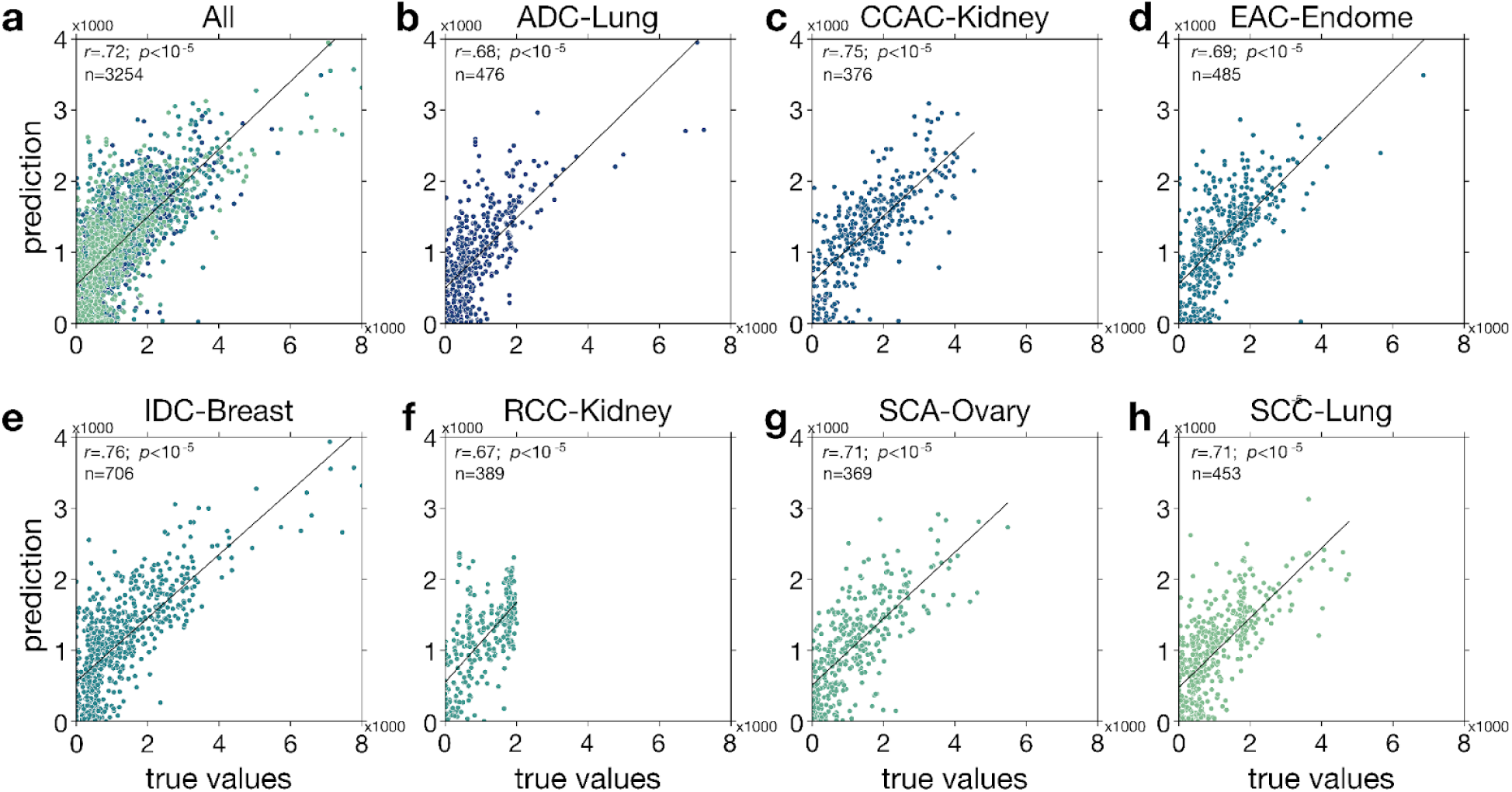
Foundational model predicts patient outcomes. Correlation between predicted and true overall survival for all histologies and resection sites (**a**) and every individual histology and resection site (**b-h**). All correlations are significant (Pearson’s correlation: *p* < 0. 05/16 ∼0. 0031, adjusted p-values for multiple comparisons).

To further investigate the consistency of model performance within different patient subgroups, we stratified the data by resection site and histology (see **Table 1**.) and computed the correlation between predicted and true values for each subset (**Figure 1.b-h**). Pearson correlations ranging from 0.67-0.76 were observed, all of which were significant (p<0.00001).

These results collectively underscore the efficacy of our fine-tuned Geneformer model in predicting clinically relevant outcomes from bulk tumor gene expression data. The strong correlations observed between predicted and true values, coupled with the consistent performance across different patient subgroups, suggest that this methodology has the potential to serve as a valuable tool for prognostication and treatment decision-making in different cancer types.

### Patient Stratification and Survival Analysis

To further explore the clinical implications of our model’s predictions, we first calculated the concordance index (c-index) to assess the predictive accuracy of the model for overall survival based on OSDTLF and DTD values. Following this, we employed a patient stratification approach, categorizing patients into three distinct risk groups—nonresponder (NR), moderate responder (M), and responder (R) tertiles—according to their predicted outcomes. This stratification allowed us to investigate the association between the model’s predicted risk categories and actual patient survival.

For the full unstratified population, the C-index was 0.77, indicating that the model correctly distinguished between patients with different survival risks 77% of the time, demonstrating a good level of concordance between predicted risk scores and actual outcomes. In order to assess survival differences for a stratified cohort, Kaplan-Meier survival curves were generated for each tertile (**Figure 2.a**). The log-rank test was employed to statistically compare the survival distributions between the three groups (**Supplementary. Table 1**). The results of the log-rank test for OS revealed highly significant differences in survival between all pairwise comparisons **Figure 2.a** ; NR vs. R: *χ*^2^ = 446 . 3, *p* < 0 . 0001 NR vs. M: *χ*^2^ = 139 . 6, *p* < 0 . 0001; R vs. M: *χ*^2^ = 184 . 0, *p* < 0 . 0001). These findings demonstrate a clear separation of survival curves, with patients in the responder tertile (R) exhibiting significantly longer OS compared to those in the moderate responder (M) and nonresponder (R) tertiles. Similarly, patients in the moderate responder tertile displayed significantly longer OS than those in the nonresponder tertile.

**Figure 2.**
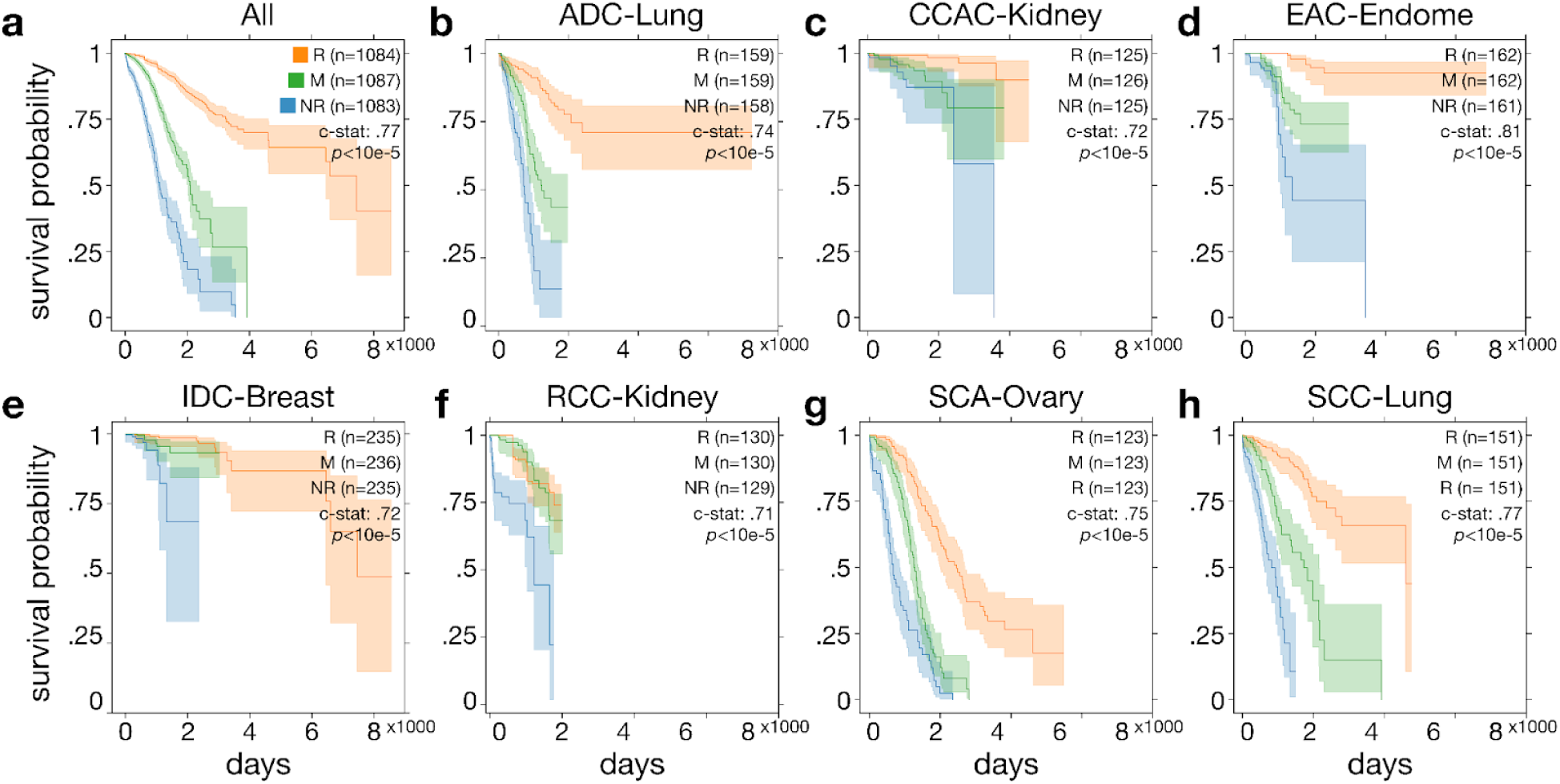
Model can stratify patients. Plots show Kaplan-Meier survival curve based on model prediction and true labels using top (orange, Responder), middle (green) and bottom (blue, NonResponder) predicted thirds of population for all histologies and resection sites (**a**) and every individual histology and cancer type (**b-h**). Curves illustrate the estimated survival probability over time, with shaded 95% confidence intervals. The curve was generated based on event duration data, with error bars representing the variability in the survival estimate. A log-rank test was performed to assess differences between the survival distributions of these subgroups (c-statistics and p-value are significant. *p* < 0. 05/16 ∼0. 0031).

To examine the consistency of these survival patterns across different patient subgroups, we calculated C-index and performed stratified survival analyses based on resection site and histology (**Figure 2.b-h, Supplementary Table 1**). While the specific survival patterns varied across subgroups, the overall trend of decreasing survival with increasing predicted risk remained largely consistent, suggesting the generalizability of our model across a variety of cancer types.

These findings collectively highlight the potential clinical utility of our model in stratifying patients into distinct risk categories based on their predicted OS. The significant differences in survival observed between the tertiles underscore the model’s ability to identify patients at high risk of adverse outcomes, potentially enabling more targeted and personalized treatment strategies.

### Stage Analysis and its Impact on Model Predictions

To further understand the influence of tumor stage on our model’s predictions and its relationship with actual patient outcomes, we performed a comprehensive stage analysis. This analysis aimed to evaluate how tumor stage affects both OS and their interaction with the predicted risk categories.

#### Stage-Based Kruskal-Wallis ANOVA

A two-way Kruskal-Wallis ANOVA was conducted to examine the effects of stage and true outcome categories (R, M, and NR tertiles based on true OS values) on the actual OS.

The analysis revealed a significant main effect of OS category (*F* = 228. 3,*p* < 0. 0001), indicating that the true OS significantly differed between the three tertiles, as expected. However, there was no significant main effect of stage (*F* = 0. 07, *p* = 0. 99) or interaction between stage and OS category (*F* = 1. 41, *p* = 0. 97), suggesting that tumor stage did not significantly influence the true OS or its relationship with the risk categories.

#### Stage-Based Spearman’s Rank Correlation

To further quantify the relationship between tumor stage and the true labels, we calculated Spearman’s rank correlation coefficient. We found a weak negative correlation (**Figure SI-1.a**; *r* =− 0. 089, *p* =< 0. 00001), suggesting a slight tendency for the OS to decrease with increasing stages. *Predicted Categories Analysis*. We then repeated the two-way Kruskal-Wallis ANOVA using the predicted categories instead of the true categories to assess the interaction between stage and the model’s predicted risk groups.

The analysis revealed a significant main effect of both stage (*F* = 15. 61, *p* = 0. 0014) and predicted OS category (*F* = 147. 17, *p* < 0. 0001). However, the interaction between stage and predicted OS category was not significant (*F* = 5. 09, *p* = 0. 53). This suggests that while both stage and the model’s predictions independently influenced OS, their combined effect was not significant.

#### Stage-Based Spearman’s Rank Correlation for Predicted Labels

Spearman’s rank correlation was also calculated between tumor stage and the predicted labels. A weak negative correlation was observed (**Figure SI-1.b**; *r* =− 0. 04, *p* = 0. 043). This suggests a subtle tendency for the model’s predicted risk to increase with advancing stage.

#### Comparison of Correlation Coefficients

We compared the correlation coefficients between “stage vs true” and “stage vs predicted” categories using Fisher’s r-to-z transformation. The difference was not statistically significant *p* = 0. 079);. This suggests that the model’s ability to capture the relationship between stage and outcome is comparable to the actual relationship observed in the data.

#### Stratification Analysis by Stage

To further investigate the potential impact of stage on the model’s stratification ability, we compared the predictive accuracy of the model across different stages using the c-index. We conducted this analysis for the entire dataset as well as within specific subgroups defined by resection site and histology, where stage information was available, aiming to assess whether the model’s stratification power is consistent or varies with tumor stage (**Figure 3.a-g**).For the entire dataset using OS as endpoint, (**Figure 3a**), C-index values were .79, .78, .77, and .68 for stages 1, 2, 3, and 4, respectively. Stratifying the population by resection site and histology revealed significant differences in c-index values across most stages (see **Figure 3.b-g** for details). These results indicate consistently high high model performance within specific cancer types and stages.

**Figure 3.**
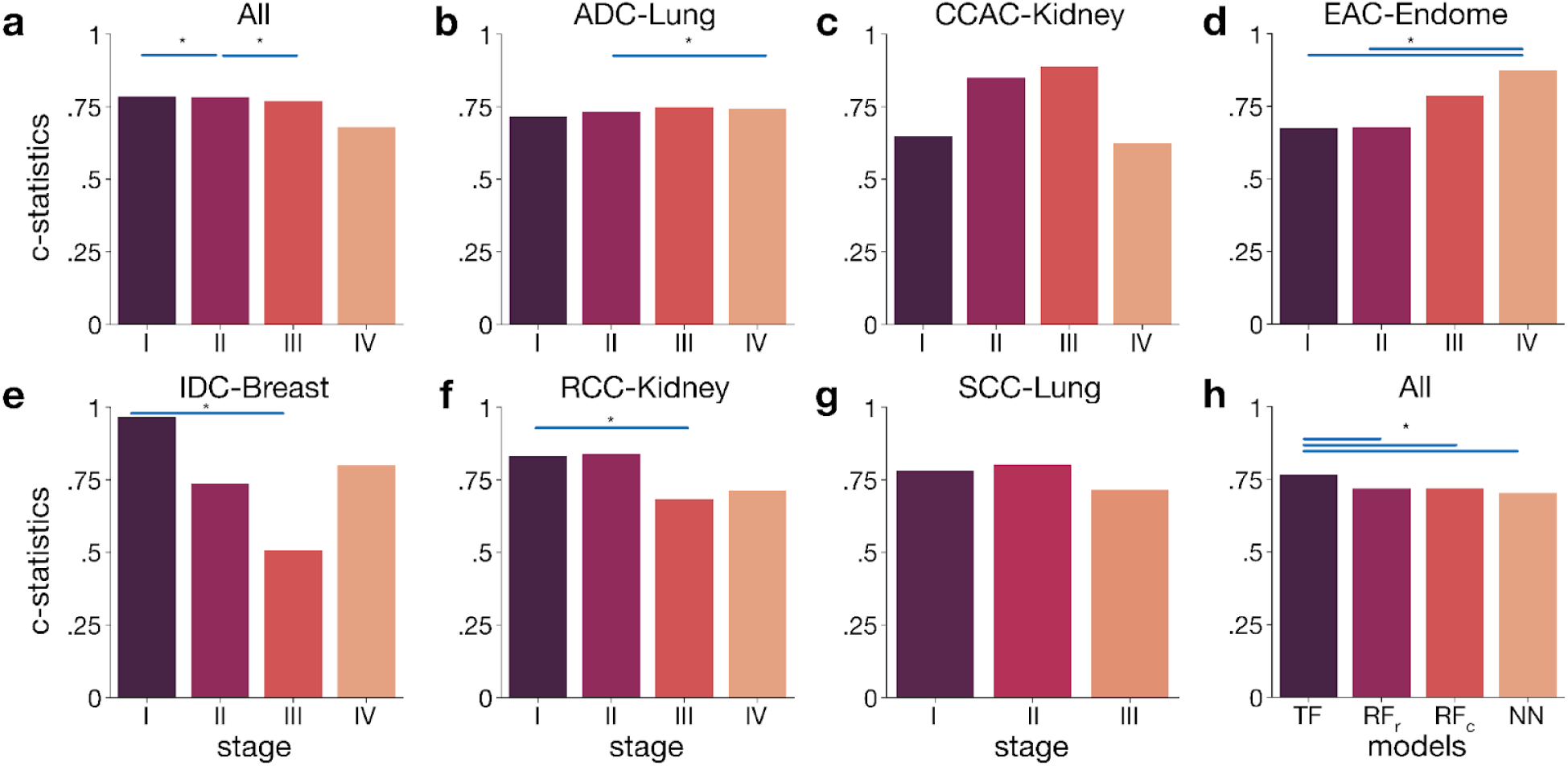
(**a-g**) **Model can stratify patients regardless of stage.** Plots show c-statistics of logrank test performed to assess differences between the survival distributions of subgroups (see **Supplementary Table 1** test-statistics of logrank test) between different stages for overall survival for all histology cites and cancer types (**a**) and every single cite and type (**b-g**). **Gene transformer performs stratify patients better than alternative models.** Plot show c-statistics of logrank test performed to assess differences between the survival distributions of subgroups between different models (TF: Gene Transformer, RF_R_: Random Forest rank, RF_c_: Random Forest count, NN: Neural Network) for overall survival (**p**) for all histology cites and cancer types. Histology sites and cancer types are shown at the top of each sub-column. Asterisks show significant differences between c-statistics between stages (*p* < 0. 05/12; Bonferroni correction).

### Model Comparison and Performance Benchmarking

Having established the prognostic potential of our Geneformer-based model, we sought to further contextualize its performance by comparing it against established machine learning approaches. We implemented two widely-used machine learning models - Random Forest (RF) and Neural Network (NN) - and evaluated their ability to stratify patients into clinically meaningful risk groups. This comparative analysis aimed to shed light on the relative strengths and weaknesses of different modeling paradigms in the context of survival outcome prediction from bulk tumor gene expression data.

For the Random Forest model, we explored two distinct feature encoding strategies: one based on gene ranking (*RF_r_*), mirroring the approach used in our Geneformer model, and another based on raw gene counts (*RF_c_*). This allowed us to assess the impact of feature encoding on the performance of the Random Forest model. The Neural Network model was implemented with a standard architecture commonly used for regression tasks.

*Comparative Analysis*. To rigorously compare the stratification abilities of the different models, we performed log-rank tests to assess the differences in survival distributions between patient subgroups stratified by each model. Additionally, we computed the correlation coefficients between the true and predicted labels for each model to provide a quantitative measure of their predictive accuracy. Our Geneformer model outperforms other models in this task, as evidenced by the higher correlation coefficients between true and predicted labels (see **Figures 1a** and **SI-2**).

Our hypothesis was that the stratification ability would vary between models, with the potential for the Geneformer-based model to outperform the traditional machine learning approaches due to its pre-training on a vast single-cell RNA-seq dataset and its ability to capture complex gene expression patterns.

We compared the stratification abilities of different models using log-rank tests to assess differences in survival distributions between subgroups stratified by each model (**Figure 3.h**). The results highlight the superior stratification performance of our Geneformer model. The log-rank tests, comparing the survival distributions across different models, demonstrate the significant outperformance of our model in this task.

## Conclusion

In this study, we successfully leveraged the power of Geneformer, a state-of-the-art transformer model pre-trained on single-cell RNA-seq data, to predict OS outcome from bulk tumor gene expression data. By adapting Geneformer to the unique challenges of bulk tumor data analysis and implementing a rank-value encoding scheme, we developed a predictive model that demonstrated strong correlations with patient outcomes. Furthermore, our model exhibited consistent performance across different patient subgroups and tumor stages, highlighting its potential for broad clinical applicability.

The superior performance of our transformer-based model compared to traditional machine learning approaches underscores the advantages of leveraging pre-trained foundational models in the context of complex biological data analysis. The model’s ability to capture intricate gene expression patterns and its adaptability to diverse clinical contexts positions it as a promising tool for prognostication and treatment decision-making in cancer research.

However, we acknowledge that this work represents a stepping stone in the ongoing pursuit of more accurate and clinically relevant predictive models. Future research should explore the development of even more sophisticated gene transformer architectures, potentially incorporating multi-omics data and leveraging larger, more diverse training datasets. Additionally, further validation of our models on external, clinically annotated datasets, including those from commercial sources, is warranted to ensure their robustness and generalizability.

By continuing to refine and expand upon these foundational approaches, we can strive towards a future where precision medicine is guided by powerful predictive models, ultimately improving patient outcomes and transforming the landscape of cancer care.

## Methods

### Data Acquisition and Preprocessing

This subsection details the acquisition and preprocessing steps applied to the data utilized in this study.

### Data Sources

The Cancer Genome Atlas (TCGA) program, established by the National Cancer Institute (NCI) and National Human Genome Research Institute (NHGRI), provides a comprehensive collection of human cancer genomic and clinical data [8]. We downloaded gene expression (RNA-Seq) and clinical data for [cancer type] from the TCGA Data Portal (https://portal.gdc.cancer.gov/).

### Data Preprocessing

Several preprocessing steps were performed to ensure the quality and consistency of the data for downstream analysis:

#### Filtering

Genes with low expression (counts per million [CPM] < 10) were excluded to minimize noise. The chosen threshold can be determined based on the specific cancer type and data distribution.

#### Normalization

Gene expression data was normalized using voom transformation to account for technical variations and sequencing bias .

### Clinical Data Integration

Clinical data downloaded from TCGA, including patient demographics, disease stage, and overall survival (OS), was integrated with the preprocessed gene expression data. This integration enables exploration of relationships between gene expression profiles and clinical outcomes.

### Rank Value Encoding

Following preprocessing, gene expression data was further encoded using a rank value encoding method inspired by [7]. This approach prioritizes genes that distinguish cell state by ranking them based on their expression within each cell normalized by their expression across the entire dataset.

Here, we leverage the pre-built tokenizer module provided by the authors, which streamlines the ranking and normalization process based on a reference dataset (Genecorpus-30M) [7]. This method offers several advantages:

*Prioritizes informative genes*: Genes with high expression variability across cells are ranked higher, emphasizing their role in defining cell state.

*Reduces noise*: Housekeeping genes with ubiquitous expression are down-ranked, minimizing their impact on downstream analysis.

*Robustness*: Ranking is less susceptible to technical artifacts compared to absolute transcript count values.

The tokenizer module ensures consistent normalization across datasets, facilitating model generalizability.

### Model Architecture and Fine-tuning

We employed the pre-trained Geneformer transformer model [7] as the foundation for our downstream tasks. Geneformer, originally trained on a massive single-cell RNA-seq dataset (Genecorpus-30M), utilizes six transformer encoder units, each comprising a self-attention layer and a feed-forward neural network layer [7]. Key architectural parameters include an input size of 2,048, an embedding dimension of 256, four attention heads per layer, and a feed-forward size of 512 [7]. The model employs full dense self-attention to maximize the context window during processing.

To adapt Geneformer to our specific prediction goals (DTLF and DTD), we implemented a two-step fine-tuning process. First, we extended the pre-trained Geneformer architecture by adding a seventh transformer layer. The weights of this additional layer were initially trained in an autoencoder-like fashion, allowing the model to further refine its representation of the input gene expression data. Subsequently, we appended a task-specific fine-tuning layer and fine-tuned the entire model on the TCGA data to predict DTLF and DTD.

For fine-tuning, we utilized all available data points, irrespective of cancer type or histology, to leverage the full diversity of the dataset. We employed a 10-fold cross-validation strategy, training the model on 90% of the data and evaluating its performance on the remaining 10% in each fold. This process was repeated ten times to ensure that predictions were generated for the entire dataset. While we retained the fine-tuning hyperparameters as described by Theodoris et al. (2023) [7] for a controlled comparison, future work may explore the impact of hyperparameter optimization on model performance for our specific prediction tasks.

### Benchmark Models and Evaluation

To provide a comparative assessment of our Geneformer-based approach, we implemented two widely-used machine learning models:

Random Forest (RF) and Neural Network (NN). Both models were trained and evaluated using a similar 10-fold cross-validation strategy as described for Geneformer.

For the Random Forest model, we explored two distinct feature encoding strategies: one utilizing the ranked gene expression values (*RF_r_*), aligning with the input format of Geneformer, and another employing raw gene counts (*RF_c_*). This allowed us to assess the impact of feature encoding on the performance of the RF model.

Given the high dimensionality of the gene expression data, we applied Recursive Feature Elimination (RFE), a simple yet effective feature selection method, to reduce the number of input genes to 100 for each of the benchmark models (*RF*_*r*_, *RF*_*c*_, and *NN*). This step aimed to enhance computational efficiency and mitigate the potential for overfitting. For the implementation of RF and NN, we leveraged readily available functionalities within the TensorFlow framework, utilizing standard architectures commonly employed for regression tasks.

### Survival Analysis

Survival analysis was conducted to evaluate the association between predicted and actual patient overall survival. Kaplan-Meier curves were generated to visualize the survival probabilities over time for each risk group. The log-rank test, a non-parametric statistical test, was employed to assess the significance of differences in survival distributions between the groups [9]. The log-rank test statistic and corresponding p-values were reported to quantify the statistical significance of the observed differences. Both true and predicted labels were used to stratify patients into risk groups, allowing us to compare the prognostic value of the models’ predictions against the actual clinical outcomes. Additionally, we performed stratified survival analyses based on resection site and histology to explore potential subgroup-specific effects.

### Concordance Index (C-Index) Calculation

The concordance index (c-index) [11] was calculated to assess the predictive accuracy of the model for overall survival, providing a measure of how well the model’s predicted risk scores correlate with actual survival outcomes.

### Log-Rank Test

The log-rank test is a widely used statistical method for comparing the survival distributions of two or more groups [10]. It is particularly suitable for analyzing time-to-event data, such as DTLF and DTD in our study, where the event of interest is either the last follow-up or death. The log-rank test calculates a test statistic based on the observed and expected number of events in each group at each time point. The null hypothesis of the log-rank test is that there is no difference in survival between the groups. A small p-value indicates evidence against the null hypothesis, suggesting a significant difference in survival distributions. In our study, we employed the log-rank test to compare the survival curves of patients stratified into different risk groups based on both true and predicted labels, enabling us to assess the prognostic value of our models’ predictions.

## Supplementary Information

**Supplementary Table 1.** Test-statistics of logrank test performed to assess differences between t<u>he survival distributions of subgroups</u>

| Population | Comparison | P-value | T-value |
| --- | --- | --- | --- |
| Adenocarcinoma, Lung | NR vs. R | $1.71 \times 10^{-18}$ | 76.998 |
| | NR vs. M | $4.54 \times 10^{-06}$ | 21.021 |
| | R vs. M | $8.77 \times 10^{-09}$ | 33.095 |
| Clear cell adenocarcinoma, Kidney | NR vs. R | $1.25 \times 10^{-05}$ | 19.077 |
|  | NR vs. M | 0.147 | 2.099 |
|  | R vs. M | 0.001 | 10.438 |
| Endometrioid adenocarcinoma, Endometrium | NR vs. R | $3.38 \times 10^{-16}$ | 66.569 |
|  | NR vs. M | 0.001 | 10.201 |
| | R vs. M | $1.23 \times 10^{-07}$ | 27.966 |
| Infiltrating duct carcinoma, Breast | NR vs. R | $1.94 \times 10^{-05}$ | 18.241 |
|  | NR vs. M | 0.011 | 6.471 |
|  | R vs. M | 0.0436 | 4.071 |
| Renal cell carcinoma, Kidney | NR vs. R | $1.06 \times 10^{-07}$ | 28.260 |
| | NR vs. M | $1.92 \times 10^{-07}$ | 27.107 |
|  | R vs. M | 0.553 | 0.352 |
| Serous cystadenocarcinoma, Ovary | NR vs. R | $2.97 \times 10^{-25}$ | 107.803 |
| | NR vs. M | $9.11 \times 10^{-07}$ | 24.107 |
| | R vs. M | $2.76 \times 10^{-14}$ | 57.895 |
| Squamous cell carcinoma, Lung | NR vs. R | $1.27 \times 10^{-21}$ | 91.245 |
| | NR vs. M | $1.62 \times 10^{-07}$ | 27.443 |
| | R vs. M | $1.55 \times 10^{-10}$ | 40.955 |
| All | NR vs. R | $4.52 \times 10^{-99}$ | 446.337 |
| | NR vs. M | $3.18 \times 10^{-32}$ | 139.645 |
| | R vs. M | $6.43 \times 10^{-42}$ | 184.017 |

**Figure SI-1.**
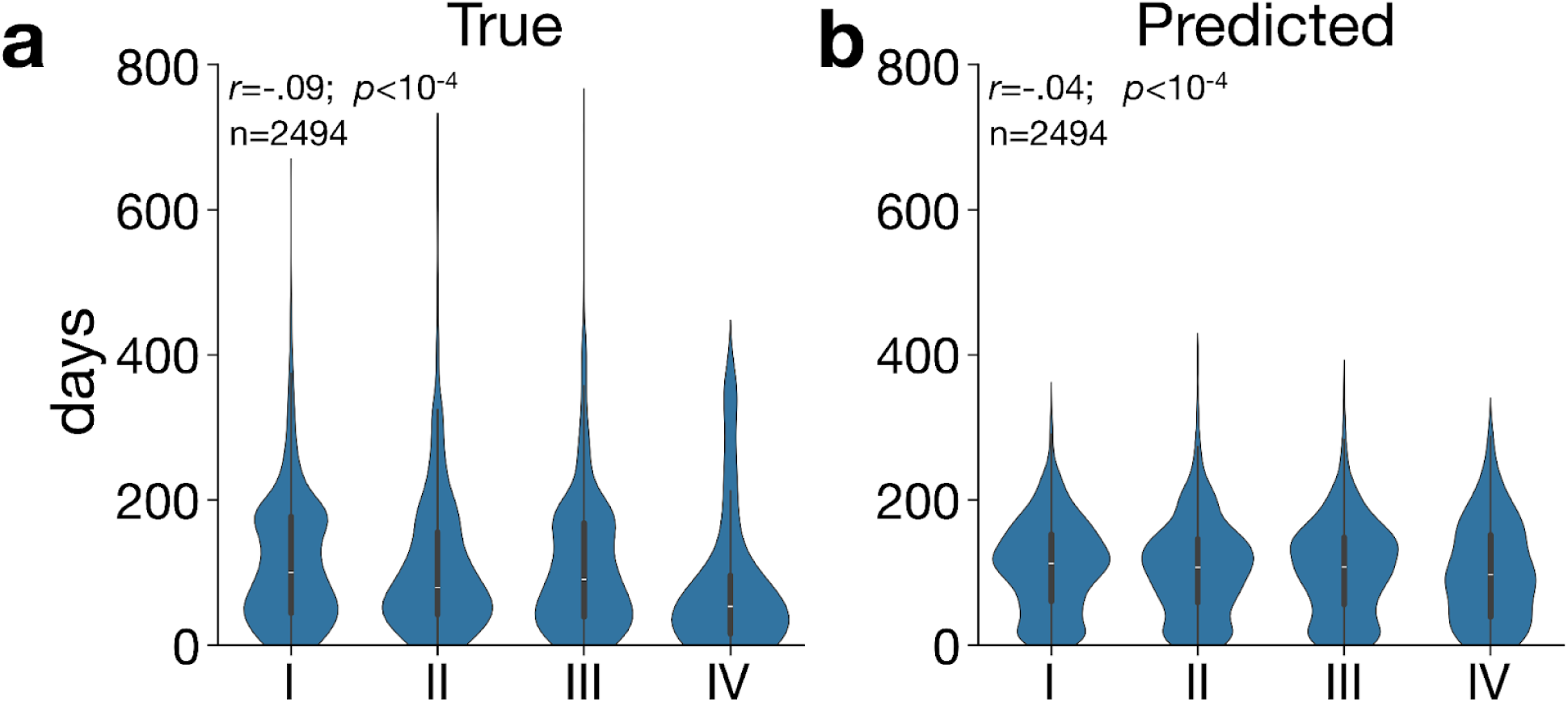
Stage has a weak negative influence on labels. Plots show violin distributions of true (**a**), and predicted (**b**) overall survival stratified by pathologic stage. All correlations are significant (Spearman’s rank correlation: *p* < 0. 05/4 ∼0. 0125, adjusted p-values for multiple comparisons).

**Figure SI-2.**
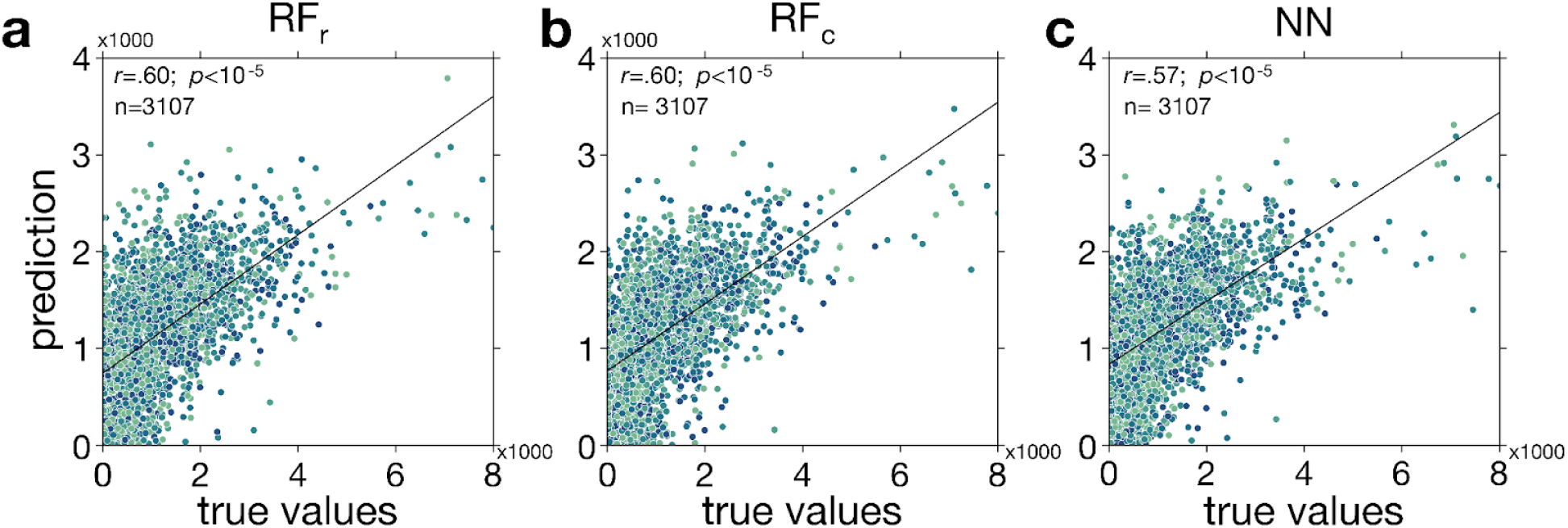
Alternative models underperform in predicting patient outcomes. Correlation between predicted and true labels for overall survival for all histology cites and different models. Column titles indicate respective model names. All correlations are significant (Pearson’s correlation: *p* < 0. 05/6 ∼0. 008, adjusted p-values for multiple comparisons).

